# CYP79B2 and CYP79B3 contribute to root branching through production of the auxin precursor indole-3-acetonitrile

**DOI:** 10.1101/2023.09.26.559630

**Authors:** Eva van Zelm, Iko T. Koevoets, A. Jessica Meyer, Kyra van der Velde, Thijs A. J. de Zeeuw, Francel Verstappen, Rens Holmer, Wouter Kohlen, Viola Willemsen, Charlotte M.M. Gommers, Christa Testerink

**Affiliations:** Laboratory of Plant Physiology, Wageningen University & Research, Wageningen, the Netherlands; Laboratory of Cell and Developmental Biology, Cluster Plant Developmental Biology, Wageningen University & Research, Wageningen, the Netherlands

## Abstract

Lateral root placement, outgrowth and density are influenced by environmental changes, including salinity stress. CYP79B2 and B3 are two cytochrome P450 enzymes previously identified as required for root architecture remodeling in salt. They produce iAOx, a metabolite that can be converted into indole glucosinolates (IGs), camalexin and indole-3-acetic acid (IAA), a type of auxin. We report here that lateral root appearance, induced by an auxin maximum in the bending zone after gravistimulation, is delayed in the absence of CYP79B2/B3. This delay traces back to a decrease in early lateral root growth after emergence, taking place before lateral roots are macroscopically visible. We measured gene transcripts and abundance of metabolites in the iAOx pathway in root segments that are forming lateral roots. Genes involved in tryptophane and IG biosynthesis were upregulated in *cyp79b2/b3* mutants, suggesting a transcriptional feedback-loop. Salt stress was found to increase the expression of genes involved in IAN biosynthesis, a precursor of both IAA and camalexin, in the root during lateral root formation. Moreover, salt increases the concentration of IAN in tissue forming lateral roots in a CYP79B2/B3 dependent manner, but these changes in IAN did not coincide with altered IAA levels. Both the reduction in lateral root density under salt and the delayed lateral root appearance in *cyp79b2/b3* knock-out mutants can be complemented by exogenous application of IAN. Our results reveal a role for the iAOx pathway in regulating the timing of lateral root appearance, allowing the modulation of lateral root density under salt stress.

## Introduction

The root system architecture of plants changes in response to abiotic stress [reviewed in (Koevoets et al., 2016; van Zelm et al., 2020)]. In dicots, the seedling root system includes both a primary (embryonic) root and lateral roots arising from primary root tissue. The formation of new lateral roots follows various stages [reviewed in (Péret et al., 2009; Du and Scheres, 2018)]. First, the formation of a new lateral root is initiated; a process in which a lateral root founder cell forms a lateral root primordium. This primordium goes through seven stages of development to form an emerged lateral root (Malamy and Benfey, 1997). Altering root growth rates and lateral root formation changes the root system in response to abiotic stresses [reviewed in (Koevoets et al., 2016)].

During salt stress, different strategies of altered root branching, and modulation of main– and lateral root length were identified in *Arabidopsis thaliana* accessions (Julkowska et al., 2014). A genome wide association study (GWAS), aimed at identifying genes underlying root plasticity during salt stress, identified *CYTOCHROME P450 79 B2 (CYP79B2)* as a candidate gene to contribute to average lateral root length per main root length under salt stress (Julkowska et al., 2017). Under salt stress, the expression of *CYP79B2* in individual accessions positively correlated with both average lateral root length per main root length and with lateral root density. Moreover, the double mutant *cyp79b2/b3* showed reduced average lateral root length and lateral root density under high salt stress. However, it remains unknown how CYP79B2 and CYP79B3 influence lateral root branching or elongation.

CYP79B2 and CYP79B3 are cytochrome P450 enzymes that convert tryptophane to indole-3-acetaldoxime (iAOx) at the start of the iAOx biosynthesis pathway (Hull et al., 2000; Mikkelsen et al., 2000) (Figure 1A). The major final products of the iAOx pathway are indole glucosinolates (IGs), camalexin and indole-3-acetic acid (IAA), a type of auxin. Conversion of iAOx to IAA is realized via the two intermediates indole-3-acetamide (IAM) and indole-3-acetonitrile (IAN). Within whole seedlings, radioactive labelling of iAOx showed incorporation into IAM, IAN and IAA. Moreover, IAN was not detectable in *cyp79b2/b3* mutants while IAM levels were greatly reduced (Sugawara et al., 2009). In contrast, IAA levels were not affected in *cyp79b2/b3* mutants under control conditions, but were reduced in *cyp79b2/b3* mutants at high temperatures in seedlings (Zhao et al., 2002; Sugawara et al., 2009). In control conditions, the biosynthesis pathway mediated by TRYPTOPHAN AMINOTRANSFERASE OF ARABIDOPSIS (TAA) and YUCCA (YUC) enzymes is considered the major route of IAA production from tryptophane, a pathway that is independent of CYP79B2/B3 action (Stepanova et al., 2011; Won et al., 2011).

**Figure 1.**
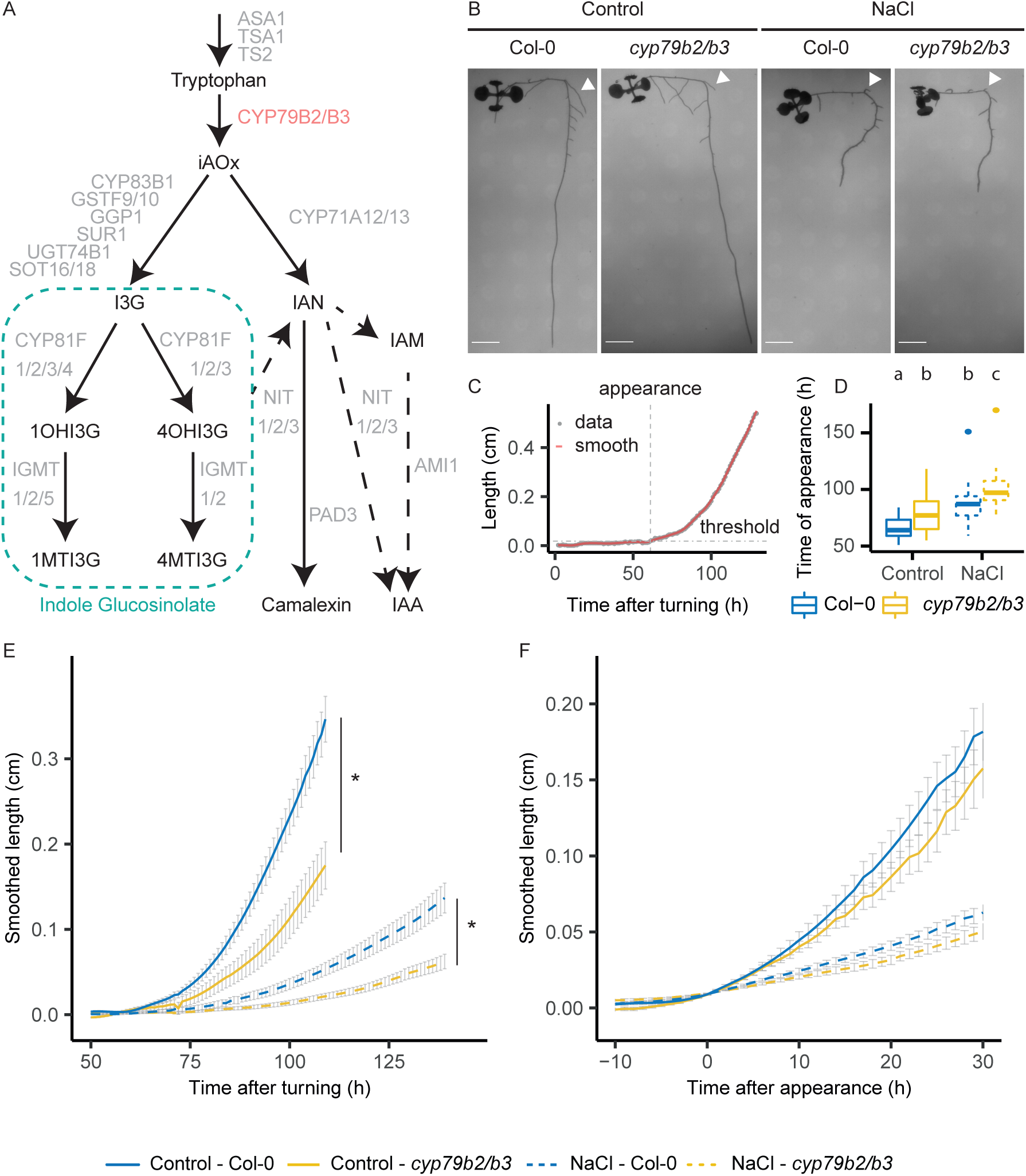
The *cyp79b2/b3* mutant shows a delay in lateral root appearance after gravistimulation. A) Simplified overview of enzymes involved in the iAOx pathway. ANTHRANILATE SYNTHASE ALPHA SUBUNIT 1 (ASA1), TRYPTOPHAN SYNTHASE ALPHA CHAIN (TSA1) and TRYPTOPHAN SYNTHASE BETA-SUBUNIT 2 (TSB2) are involved in the production of tryptophane. CYP79B2/B3 convert tryptophane to iAOx. CYP83B1, GLUTATHIONE S-TRANSFERASE PHI 9 (GSTF9), GSTF10, GAMMA-GLUTAMYL PEPTIDASE 1 (GGP1), SUPERROOT 1 (SUR1), UDP-GLUCOSYL TRANSFERASE 74B1 (UGT74B1), SULFOTRANSFERASE 16 (SOT16) and SOT18 convert iAOx to I3G and thereby form the core IG structure. Subsequently, side chain modification of IGs are performed by different members of the CYP81 and INDOLE GLUCOSINOLATE O-METHYLTRANSFERASE (IGMT) family. Alternatively, iAOx can be converted to IAN by CYP71A12 and CYP71A13. Subsequently, PAD3 can convert IAN to camalexin. The NITs possibly convert IAN to IAA, but might also be involved in converting IG to IAN. AMI1 is involved in converting IAM to IAA. [this pathway is adapted from (Lehmann et al., 2017; Vik et al., 2018; Crane et al., 2019; Harun et al., 2020)]. Black text = metabolites, grey text = enzymes, red text = CYP79B2/B3, turquoise box = indole glucosinolates, arrows = one or more enzymatic steps, dashed arrows = hypothesized enzymatic steps. B – F) To induce the synchronized formation of a lateral root in the bending zone, plants were turned 90 degrees and after 3 hours transferred to salt (125mM NaCl) or control plates. The length of the lateral root in the bending zone was imaged every 20 min. B) Example images of lateral roots formed in the bending zone of Col-0 and *cyp79b2/b3* mutant in control conditions (5 days after turning) and under salt stress (7 days after turning). Arrow head = lateral root in bending zone, scale bar = 0.5 cm. C) Example of the length increase over time for a root in the bending zone. The threshold was defined as the average length in the first 60 hours after salt stress + 0.01 cm. A lowess smooth was applied to determine the moment of lateral root appearance (time at which the the lowess smooth > threshold). D) Timing of appearance of the lateral root (in hours after transfer) in the bending zone of Col-0 and *cyp79b2/b3* seedlings in control conditions and under salt (125mM NaCl) (statistically different groups are indicated with letters, Wilcoxon test, p < 0.05, n >= 15, pooled from 3 independent replica’s). E) Average smoothed length of the lateral root in the bending zone after transfer to salt (125mM NaCl) or control plates of Col-0 and *cyp79b2/b3* seedlings (statistical differences between genotypes per treatment shown by *, multiple Wilcoxon tests per timepoint with BH adjustment for multiple testing, in control p < 0.05 from 70 hour onwards, in NaCl p < 0.05 from 94 hour onwards, n >= 15, pooled from 3 independent experiments). F) Length of the lateral root in the bending zone after lateral root appearance under salt (125mM NaCl) or control conditions for Col-0 and *cyp79b2/b3* seedlings (no statistical differences between genotypes, multiple Wilcoxon tests per timepoint with BH adjustment for multiple testing, n >= 15, pooled from 3 independent replicates).

Several enzymes are necessary to convert iAOx into IAA (Figure 1A). The conversion of iAOx into IAN is facilitated by CYP71A12 and CYP71A13 (Böttcher et al., 2009; Müller et al., 2015). Subsequently, NITRILASES (NITs) are thought to convert IAN to IAA, but NITs might also play a role in converting IGs to IAN (Lehmann et al., 2017). AMIDASE 1 (AMI1) ultimately converts IAM to IAA (Neu et al., 2007). Under salt stress, IAA, IAN and IAM levels were shown to be increased in whole seedlings after 2 weeks (Cackett et al., 2022), but remain unchanged or are even decreased after 3 hours of salt stress (Smolko et al., 2021). IAA inhibits main root growth and promotes lateral root initiation and outgrowth (Ivanchenko et al., 2010; Du and Scheres, 2018). The application of IAN or IAM also reduced main root length and this response depended in part on NITs and AMI1 (Lehmann et al., 2017; Pérez-Alonso et al., 2021). NIT1 overexpression increased lateral root density and reduced lateral root length, while *nit1-3* mutants showed a reduced lateral root density (Lehmann et al., 2017). Thus, iAOx can be converted to IAA, and both IAA and intermediates IAN and IAM influence root growth and development.

Camalexin and IGs are known for their role in defense against herbivores and pests [reviewed in (Halkier and Gershenzon, 2006; Ahuja et al., 2012)]. IGs are synthesized from iAOx, and *cyp79b2/b3* mutants have been documented to lack IGs (Zhao et al., 2002; Halkier and Gershenzon, 2006). Furthermore, IG abundance was shown to increase upon salt stress (Chen et al., 2017). Camalexin can be formed from IAN by PHYTOALEXIN DEFICIENT 3 (PAD3) (Glawischnig et al., 2004; Nafisi et al., 2007; Böttcher et al., 2009). Even though camalexin was mostly reported in the shoot, it was also detected in root exudates upon FLAGELLIN 22 (FLG22) treatment (Millet et al., 2010). Recently, camalexin was shown to promote lateral root development (Serrano-Ron et al., 2021). Thus, to date it remains unknown which branches of the iAOx pathway are responsible for the root phenotypes observed in *cyp79b2/b3* mutants under salt stress.

Here we show that when formation of lateral roots is synchronized in the bending zone after gravistimulation, lateral roots appear later in the *cyp79b2/b3* mutant under both salt and control conditions. This delay in appearance traces back to a delay in early lateral root growth, revealing a so far uncharacterized point of control in lateral root development. Analysis of gene expression, specifically in the region which develops a new lateral root, showed that the lack of iAOx induces the expression of IG– and tryptophane-producing enzymes, revealing a transcriptional feedback loop responding to the lack of iAOx or IGs. Salt represses the expression of IG producing enzymes and induces the expression of IAN synthesizing enzymes in the same tissue. IAN levels are increased in response to salt during lateral root emergence and although the concentration of IAN was decreased in *cyp79b2/b3* mutants, the IAA concentration remained unchanged in the mutant. Exogenous application of IAN could complement the lateral root phenotypes of the *cyp79b2/b3* mutant. Together, our results show a role for the iAOx pathway in early lateral root growth which contributes to modulation of root architecture under salt stress.

## Results

### The iAOx pathway influences early lateral root elongation after a gravistimulus

Previously, *cyp79b2/b3* mutants were shown to exhibit reduced average lateral root length and lateral root density under salt stress (Julkowska et al., 2017). We aimed to investigate if the lateral root phenotypes observed in *cyp79b2/b3* mutants are due to reduced root elongation or a delay in lateral root formation. Turning seedlings 90° stimulates a gravitropic response of the main root and induces the formation of a lateral root in the bending zone (Figure 1B). The development of the lateral root is synchronized due to the formation of an auxin maximum in the bending zone (Lucas et al., 2008; Péret et al., 2012; Voß et al., 2015). Under salt stress, the timing of lateral root appearance in a turning assay has been shown to correlate with changes in lateral root density in overall root architecture (van Zelm et al., 2023). By measuring the length of the lateral root in the bending zone every 20 min after turning the seedlings, we quantified both lateral root elongation and the timing of macroscopic lateral root appearance. The latter is defined as the time at which the lateral root length reaches a set threshold (average length of the first 60 hours + 0.01 cm) (Figure 1C). In *cyp79b2/b3* mutants, the timing of lateral root appearance was later in both control and salt conditions as compared to Col-0 (p < 0.05) (Figure 1D). The length of the lateral root in the bending zone was reduced in *cyp79b2/b3* mutants compared to Col-0 in control (p < 0.05 from 70 h after turning) and in salt conditions (p < 0.05 from 94 h after turning) (Figure 1E). However, when the length of the lateral root is plotted for the time after the lateral root appearance instead of the time after the gravistimulus, there was no statistical difference in macroscopically visible lateral root elongation between *cyp79b2/b3* mutants and Col-0 (Figure 1F, p > 0.05). This indicates that the difference in lateral root length in *cyp79b2/b3* mutants is caused by a delay in lateral root appearance rather than a reduction of lateral root elongation after appearance.

Lateral root primordia develop through a series of seven developmental stages [reviewed in (Péret et al., 2009; Du and Scheres, 2018)]. To find out if the delay in lateral root appearance in the *cyp79b2/b3* mutant is caused by a delay in the early steps of lateral root development, we scored the stages of lateral root primordia in the bending zone of gravistimulated seedlings [as described in (Voß et al., 2015)]. This revealed a delay in development of approximately 1 day under salt compared to control, but surprisingly, there were no significant differences in the distribution of stages between *cyp79b2/b3* and Col-0 seedlings at 1, 2, 3 or 4 days after turning (p > 0.05), (Figure 2A). Thus, the *cyp79b2/b3* mutant does not show defects in proceeding through the stages of lateral root primordia development. A difference in macroscopic lateral root appearance, without a difference in lateral root emergence, could be explained by reduced early lateral root growth in the *cyp79b2/b3* mutant, which is not yet visible at a macroscopic level.

**Figure 2:**
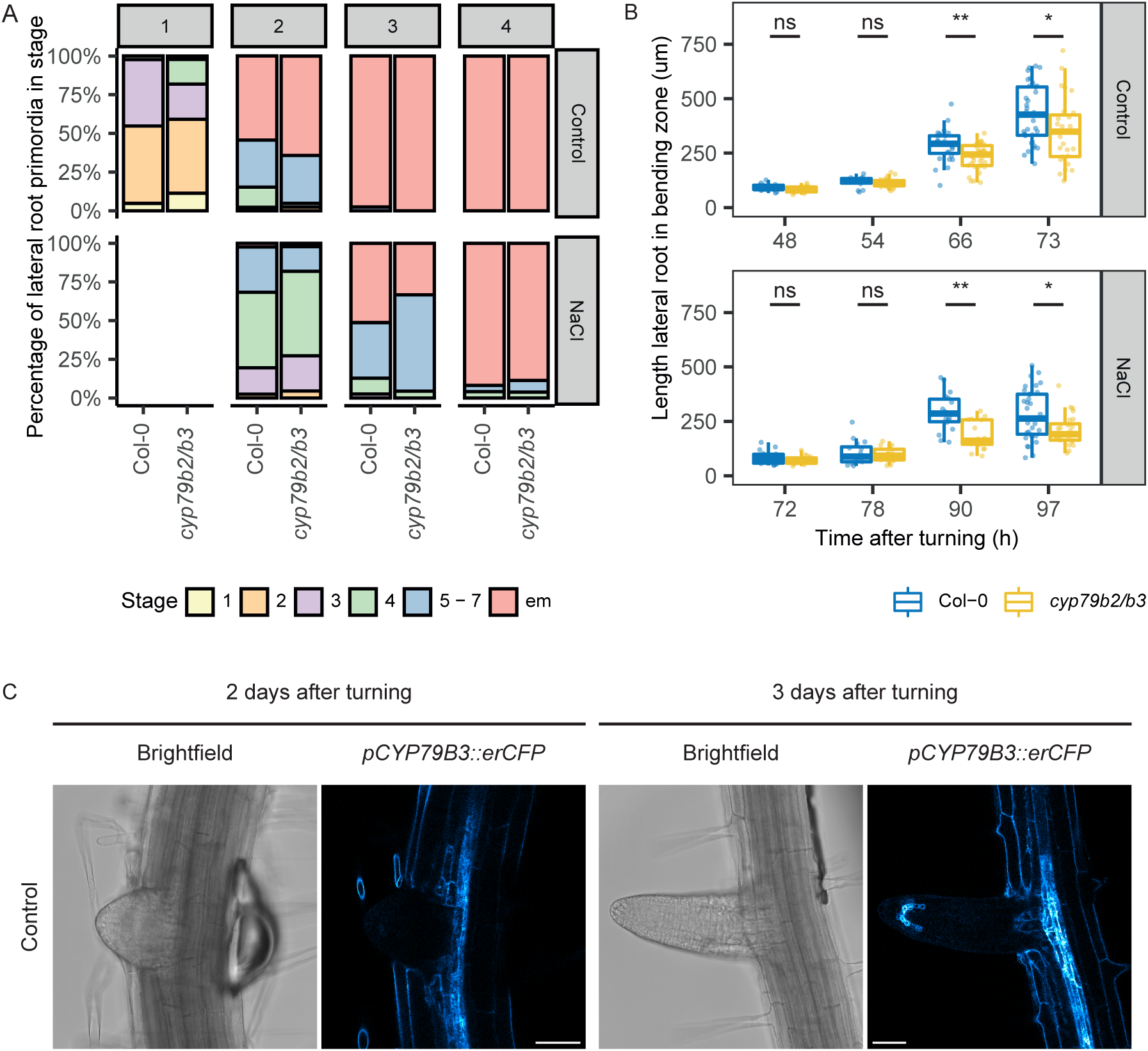
CYP79B2/B3 do not influence lateral root development but influence the early elongation of emerged lateral roots after a gravistimulus. Seedlings were turned 90 degrees to induce lateral root formation and transferred to 125 mM NaCl or control plates 3 hours after turning. A) Frequency of different stages for the lateral root in the bending zone at 1, 2, 3 or 4 days after turning (days after turning is shown on the top). No statistical differences between genotypes were found (Chi-square, n >= 39, pooled from 2 independent experiments). B) The extension of newly emerged lateral roots measured using a light microscope and quantified from the tip of the lateral root to the stele of the primary root at different timepoints after turning. Statistical differences between genotypes are shown using brackets (Wilcoxon test, ns = not significant, * < 0.05, ** < 0.01, n >= 15). C) Expression of *pCYP79B3::erCFP* in lateral roots in the bending zone of gravistimulated seedlings at 2 and 3 days after turning under control conditions. Scale bar = 50 µM.

To investigate whether the phenotype would arise between emergence of lateral root primordia and macroscopic lateral root appearance, we measured the length of newly emerged lateral roots from the tip of the lateral root to its base at the stele of the main root using a light microscope (Figure 2B). This length was quantified in the bending zone of gravistimulated seedlings after lateral root primordia emerged (48, 54, 66 and 73 h after gravistimulus for control and 72, 78, 90, and 97 h for NaCl treatment). No significant difference between the genotypes was observed at the early timepoints (48 and 54 for control and 72 and 78 for NaCl), but at the later timepoints (66 and 73 h for control and 90, and 97 h for NaCl treatment) *cyp79b2/b3* mutants had significantly shorter lateral roots compared to the wild type (p < 0.05) (Figure 2B). This supports the hypothesis that reduced lateral root length and delayed root appearance in *cyp7b2/b3* mutants (Figure 1) is caused by a reduction in lateral root elongation after emergence, but before being visible by eye, hereafter referred to as early lateral root elongation.

*CYP79B2* and *CYP79B3* are expressed in the primary and lateral root meristem and in the tissue underlying lateral root primordia, shown using a promotor-GUS reporter (Ljung et al., 2005). To investigate whether the lateral root phenotypes correlate with the expression of *CYP79B3*, we followed promotor activity of *pCYP79B3::erCFP* in newly emerged lateral roots at 2 or 3 days after turning in control conditions. A change in expression domain was observed from initially only the base of the lateral root at 2 days after turning, to both the base and the meristem of the lateral root at 3 days after turning (Figure 2C). In conclusion, *cyp79b/b3* mutants show a delay in early lateral root growth, without differences in lateral root development or lateral root elongation after appearance and this phenotype coincides with a change in CYP79B3 localization in the developing lateral root.

### Salt-induced delay in gene expression patterns coincides with delay in lateral root development

To investigate how salt and the iAOx pathway affect early lateral root elongation, we measured gene expression in the bending zones of gravistimulated Col-0 and *cyp79b2/b3* seedlings with and without salt stress by RNA sequencing at 1, 2 and 3 (control) or 2, 3 and 4 (NaCl) days after turning (Figure 3A & supplemental figure 1). Due to the 1-day developmental delay in lateral root development in salt (Figure 2A), this data can be compared either at similar timepoints or at similar lateral root developmental stages (Figure 3B, Supplemental figures 2 & 3). First, we investigated how the developmental delay influenced salt-affected gene expression in Col-0 (Figure 3, supplemental figures 2 & 3).

**Figure 3:**
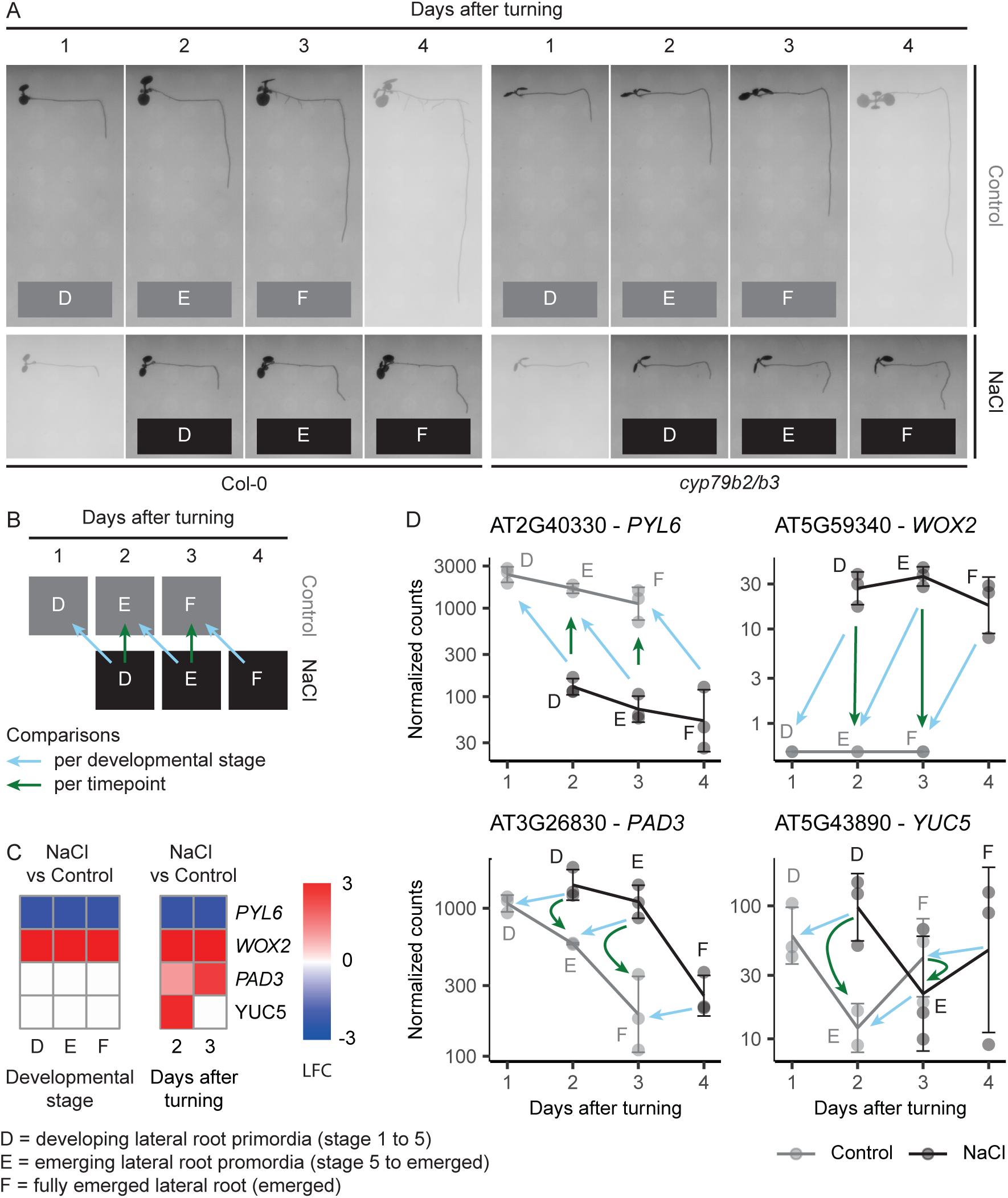
Salt-induced differential gene expression patterns during lateral root development; distinguishing the effects of delay in lateral root development from time of treatment. Seedlings were turned 90 degrees and 3 hours later transferred to 125 mM NaCl or control plates, 3 mm bending zone was dissected at 1, 2 and 3 (control) or 2, 3 and 4 (NaCl) days after turning and used for RNAseq analysis (n = 3). A) Example images of seedlings are shown at the time of dissection. Developmental stage represents the developmental stage of the lateral root in the bending zone as measured in figure 3A and are divided into 3 groups: developing lateral root primordia (D, stage 1 to 5), emerging lateral root primordia (E, stage 5 to emerged) and fully emerged lateral roots (F). B) Schematic representation of the possible comparisons of samples per developmental stage or per timepoint after turning. C) Heatmap of LFC between salt and control of *PYL6, WOX2, PAD3* and *YUC5* when quantified per developmental stage or per timepoint after turning. Only significant (p < 0.05) LFCs are shown, non-significant LFC are in white. D) Normalized gene counts of *PYL6, WOX2, PAD3* and *YUC5.* Error bar = standard deviation around mean, dots = data points, letters indicate developmental stages, arrows indicate type of comparisons. D = developing lateral roots, E = emerging lateral roots, F = fully emerged lateral roots.

The stages of lateral root development in the bending zone were divided into 3 groups (Figure 3): developing lateral roots (D, stage 1 to 5), emerging lateral roots (E, stage 5 – emerged) and fully emerged lateral roots (F) based on the stages defined in figure 2A. We identified a group of genes exhibiting differential expression in response to salt, regardless of the developmental stage or timepoint (for example *PYR1-LIKE6 (PYL6)* and *WUSCHEL RELATED HOMEOBOX2 (WOX2)* in figures 3C & 3D, group IV and V in supplemental figures 2 & 3). However, other genes are differentially expressed only when data is compared per timepoint but not when data is compared per developmental stage (for example *PAD3* and *YUC5* in figure 3C & 3D, group II and III in supplemental figures 2 & 3). Some of these genes show similar changes in expression between the same developmental stage in control and salt conditions, *i.e.* their expression pattern is delayed by a day in salt compared to control and thereby coincides with the one day developmental delay in lateral root development (for example *PAD3* and *YUC5* in Figure 3D, group III in supplemental figure 2 & 3). Thus, the differential expression of these genes either influences or is dependent on the salt-induced developmental delay in lateral root formation.

We further investigated how many and what type of genes show an expression pattern that is influenced by the salt-induced developmental delay in lateral root formation. Of the differentially expressed genes in salt stress (salt DEGs), 27% was influenced by the developmental delay at 2 days after turning (Supplemental figure 2, group III) and 11% at 3 days after turning (Supplemental figure 3, group III). At 2 days after turning, this group of salt DEGs was enriched in ribosome biogenesis, rRNA processing and fatty acid related GO terms (Supplemental dataset 1). Interestingly, fatty acid metabolism has recently been shown to be important for lateral root development (Trinh et al., 2019; Bellande et al., 2022). At 3 days after turning, this group of salt DEGs was enriched in cell growth and cell wall organization related GO terms (Supplemental dataset 1). The cell wall has been shown to play multiple roles in lateral root development, especially during emergence, where the lateral root primordium breaks through the overlaying cortex and endodermal cells [(Zhu et al., 2019), and reviewed in (Stoeckle et al., 2018)].

*PAD3*, one of the examples mentioned above as a gene influenced by the salt-induced developmental delay is part of the iAOx pathway. PAD3 converts IAN to camalexin and is downregulated during later stages of lateral root development and this downregulation is delayed in salt (Figure 3D). Interestingly, when solely looking at the DEGs per timepoint, one could conclude that *PAD3* is upregulated by salt (Figure 3C), but by cross-comparing lateral root stage-dependent expression, we now show that this downregulation is either dependent on or responsible for the salt-induced developmental delay. Our approach underlines the importance of regarding a delay in development due to stress when analyzing transcriptomics data.

### Tryptophane and IG biosynthesis genes are upregulated in cyp79b2/b3 mutants

To investigate the downstream transcriptional changes dependent on CYP79B2/B3, we identified DEGs (p < 0.05, absolute LFC > 0.5) for the *cyp79b2/b3* mutant compared to Col-0 in control conditions and for the interaction between genotype and treatment at different developmental stages (D, E, F) or at different timepoints after turning (Supplemental figure 1, supplemental dataset 2). Per timepoint or developmental stage, less than 7 genes showed a significant interaction effect, indicating that the *cyp79b2/b3* mutation has little effect on the transcriptional changes under salt stress in the bending zone (Supplemental figures 1C & 1D, supplemental dataset 2). Surprisingly, *CYP79B3* was found to be upregulated in the *cyp79b2/b3* mutant (Figure 4). However, the t-DNA insertion in *CYP79B3* is in the second exon and aligning the reads to *CYP79B3* showed that the upregulated transcripts were truncated and mostly align to the first exon (Supplemental figure 4A).

**Figure 4:**
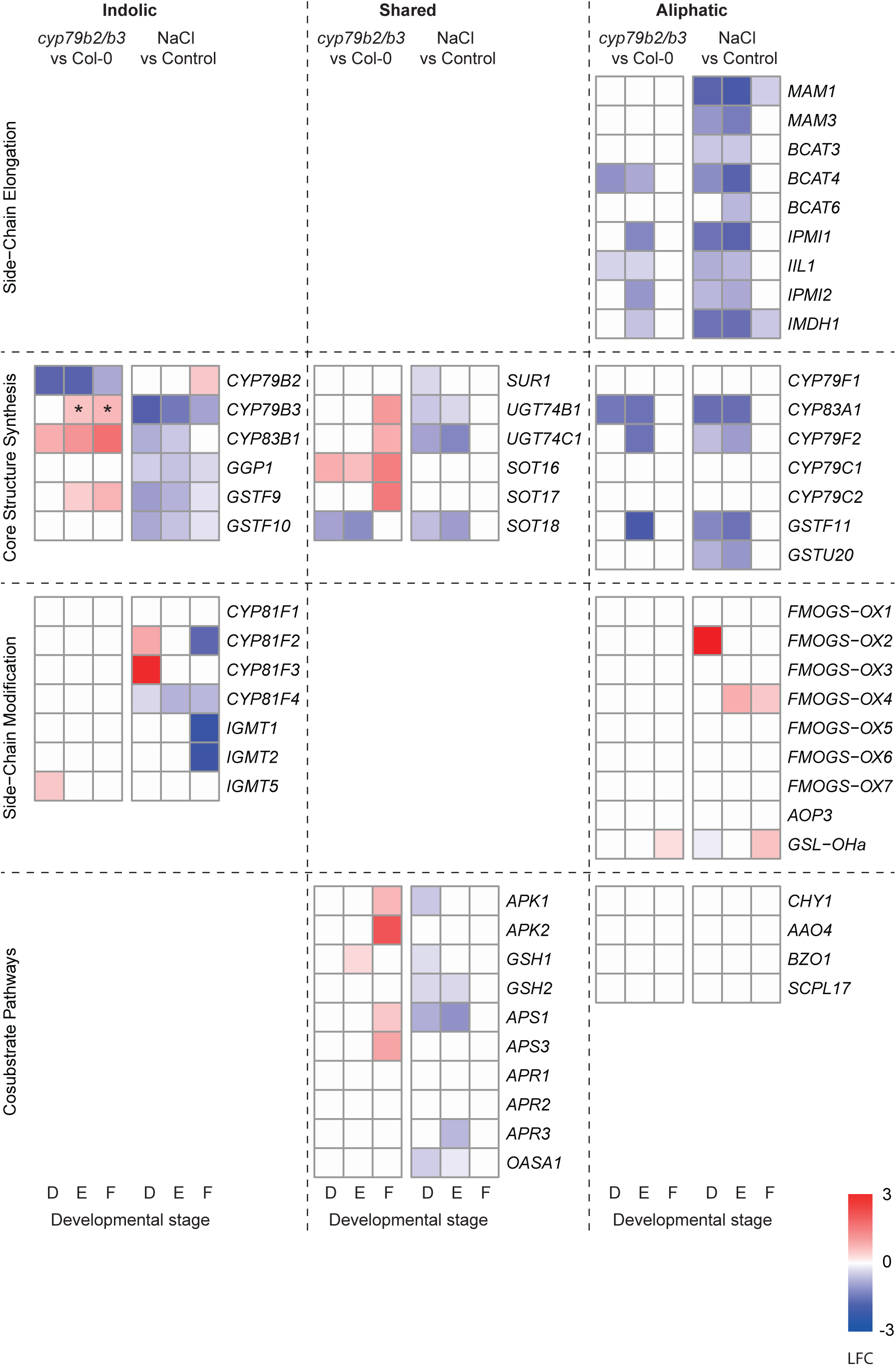
In *cyp79b2/b3* mutants genes in the indolic glucosinolate pathway are upregulated, while those in the aliphatic pathway are downregulated. Expression of enzymes with a known function in the glucosinolate biosynthesis pathway [based on (Harun et al., 2020)]. Heatmaps of LFC between *cyp79b2/b3* and Col-0 in control treatment and between NaCl and control treatment in Col-0 at different developmental stages (only expression changes with p < 0.05 are shown, non-significant changes are white). Genes are divided by their function in the aliphatic or indolic pathway or shared between the aliphatic and indolic pathway (shared). The functionality of genes is divided on their role in side-chain elongation, core structure synthesis, co-substrate pathways or side-chain modifications [based on (Harun et al., 2020)]. D = developing lateral roots, E = emerging lateral roots, F = fully emerged lateral roots. *a truncated *CYP79B3* transcript is upregulated in the *cyp79b2/b3* mutant (Supplemental figure 4A).

A GO term enrichment on differentially expressed genes between *cyp79b2/b3* and Col-0 at any developmental stage in the control treatment, revealed that genes involved in tryptophan metabolism are upregulated in the *cyp79b2/b3* mutant and several glucosinolate metabolism-related genes are upregulated while others are downregulated in the *cyp79b2/b3* mutant (Supplemental figure 4B, & supplemental dataset 2). Glucosinolates can be divided into different groups depending on the amino acids that they are synthesized from [reviewed in (Sønderby et al., 2010; Chhajed et al., 2020; Harun et al., 2020)]. Indolic glucosinolates are produced from tryptophane, in a CYP79B2/B3 dependent manner and are thereby part of the iAOx pathway. Aliphatic glucosinolates are produced from alanine, leucine, isoleucine, valine, glutamate, or methionine, while benzenic glucosinolates are synthesized from phenylalanine or tyrosine. Both aliphatic and benzenic glucosinolates are produced independently from CYP79B2/B3. Glucosinolates are produced by the following consecutive steps: 1) side-chain elongation of precursor amino-acids, 2) core-structure synthesis and 3) side-chain modifications [reviewed in (Sønderby et al., 2010; Chhajed et al., 2020; Harun et al., 2020)]. Furthermore, co-substrate pathways are necessary to produce glucosinolates such as the pathway producing sulfur donors necessary for the core-glucosinolate structure. For the genes that were differentially expressed in the *cyp79b2/b3* mutant, we investigated to which of these pathways they belonged based on a recent inventory of genes involved in glucosinolate biosynthesis (Harun et al., 2020).

Interestingly, all the downregulated glucosinolate biosynthesis genes in the *cyp79b2/b3* mutant act in the aliphatic pathway, except for *CYP79B2* (and possibly *CYP79B3* when regarding functional transcripts) itself (Figure 4 & supplemental figure 4A). In contrast, the upregulated glucosinolate biosynthesis genes either act in the indolic pathway, in the co-substrate pathway or are shared between the aliphatic and indolic pathway (Figure 4). This data suggests a transcriptional feedback loop that upregulates tryptophane and IG biosynthesis genes upon low levels of iAOx or IG in the *cyp79b2/b3* mutant. In contrast, glucosinolate biosynthesis genes in the aliphatic pathway are downregulated in the *cyp79b2/b3* mutant, suggesting that the indolic and aliphatic pathways are coregulated to balance glucosinolate production. Under salt stress, most of the glucosinolate biosynthesis genes are downregulated except for those responsible for side-chain modifications (Figure 4).

### IAN is produced during lateral root emergence under salt stress and can rescue the cyp79b2/b3 lateral root phenotypes

Besides the above-described upregulation of tryptophane and IG biosynthesis genes, there were no other genes of the iAOx pathway differentially expressed in *cyp79b2/b3*. However, salt does influence the expression of various enzymes in the iAOx pathway. *CYP79B2* and *CYP79B3*, representing the initial step, respond differently to salt stress (Figure 5A). *CYP79B2* is upregulated upon salt stress in fully emerged lateral roots (p < 0.05), while *CYP79B3* is downregulated in salt stress across all developmental stages (p < 0.05) (Figure 4 & 5A). Moreover, besides the earlier described differences in IG, tryptophane and camalexin producing enzymes (Figures 3 & 4), salt induces the expression of several IAN and IAA producing enzymes (Figure 1A & 5B).

**Figure 5:**
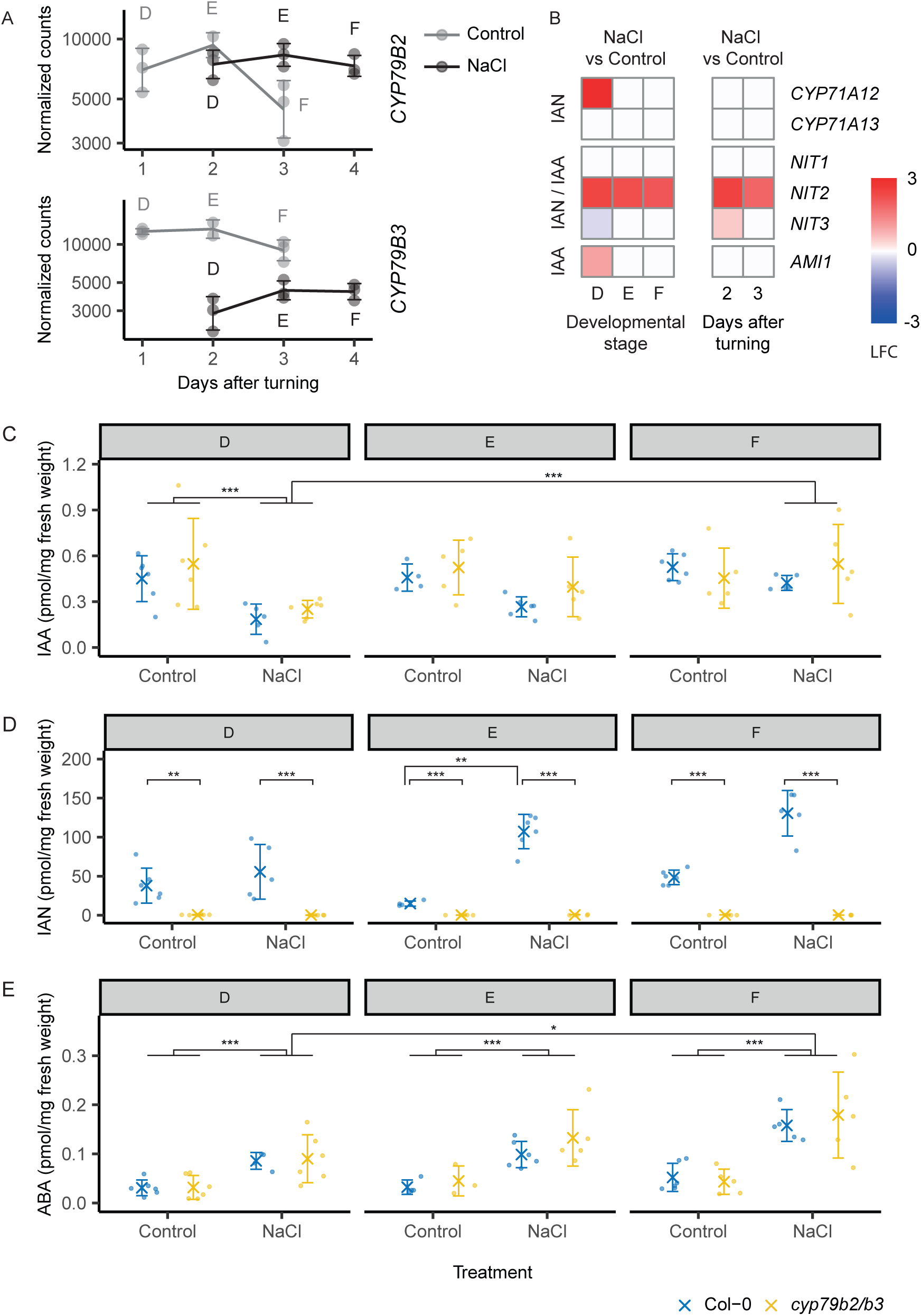
IAN concentration increases in emerging lateral roots during salt stress. A) Expression of *CYP79B2/B3* under salt stress and control conditions in the bending zone of gravistimulated seedlings. Error bar = standard deviation around mean and dots = data points. D = developing lateral roots, E = emerging lateral roots, F = fully emerged lateral roots. B) Expression of IAN and IAA producing enzymes under salt stress in the bending zone of gravistimulated seedlings. Heatmaps of LFC between NaCl and control treatment in Col-0 at different developmental stages or timepoints after turning. Only expression changes with p < 0.05 are shown, non-significant changes are white. Concentration of C) ABA, D) IAA and E) IAN in the bending zones of gravistimulated seedlings under salt and control conditions. Seedlings were transferred to 125 mM NaCl or control plates 3 hours after a gravistimulus to induce lateral root formation. Hormone concentrations were measured using UHPLC-MS/MS in in 3 mm segments of bending zones of gravistimulated seedlings. Hormone concentrations are shown relatively to the sample fresh weight. Samples were grouped per average developmental stage of the lateral root primordia: D = developing lateral roots (control: 1 day after treatment, salt: 2 days after treatment), E = emerging lateral roots (control: 2 day after treatment, salt: 3 days after treatment), F = fully emerged lateral roots (control: 3 day after treatment, salt: 4 days after treatment). X = mean, error bars = standard error and dots = datapoints. Statistical significance is indicated with brackets (3-way nonparametric factorial ANOVA, ART-C post-hoc test, * p < 0.05, ** p < 0.01, *** p < 0.001, n = 3 – 6).

As salt induced expression of IAN and IAA producing enzymes (Figure 5B) and induced a delay in *PAD3* expression (Figure 3), we quantified IAN, IAA and camalexin in developing lateral roots in the bending zone by UHPLC-MS/MS (Figure 5C-D). Camalexin was not detectable in our samples, based on a detection limit of 15 fmol (S/N >10) at injection and total recovery of the purification procedure of 46%. The concentration of IAA was not influenced by the *cyp79b2/b3* mutation, but was significantly affected by salt treatment, developmental stage, and the interaction between the two (p < 0.05). Salt decreased the concentration of IAA in early stages (D) of lateral root development, but in emerged lateral roots (stage E and F) we found no difference between salt and control treated roots. Salt treatment increased the concentration of IAA in fully emerged lateral roots (F) compared to the early stages of development (D) (p < 0.05). In contrast to IAA, IAN concentrations are close to zero in *cyp79b2/b3* mutants across all developmental stages, both in salt and control treatment (p < 0.05). Interestingly, salt increases IAN concentrations during lateral root emergence (stage E) in Col-0 seedlings (p < 0.05). This increase precedes the previously described changes in early lateral root elongation (Figure 2B) and occurs after the salt-induced upregulation of *CYP71A12* (Figure 5B), one of the enzymes that can convert iAOx to IAN (Figure 1A).

Additionally, we measured the concentration of the stress hormone abscisic acid (ABA) to confirm that we could detect effects of salt stress on hormone abundance during lateral root development and growth in small root sections (Figure 5E). Previously, salt stress was shown to increase ABA marker gene expression in lateral roots just after emergence at 4 days after treatment (Duan et al., 2013). ABA concentration was significantly influenced by salt treatment, developmental stage and the interaction between the two, but there were no differences in ABA concentration between Col-0 and *cyp79b2/b3* (p > 0.05) as expected, since it is not a product of the iAOx pathway (Figure 1A). In all stages of lateral root development, salt increased ABA concentrations in the bending zone (p < 0.05). In agreement with the measured marker gene expression (Duan et al., 2013), emerged lateral roots in salt stress had more ABA compared to lateral root primordia in early stages of development (p < 0.05).

Because of the *cyp79b2/b3-*dependent increase in IAN in response to salt treatment, we investigated whether exogenous application of IAN could complement the lateral root phenotypes of *cyp79b2/b3* mutants. IAN, but not IAM, complemented the reduced length of the lateral root in the bending zone of gravistimulated *cyp79b2/b3* mutants when compared to wild type (p < 0.05) (Figure 6A). Previous research showed a reduced lateral root density in the *cyp79b2/b3* mutant grown under salt stress (Julkowska et al., 2017). We found that also this root architecture phenotype could be complemented by the exogenous application of IAN (Figure 6). Interestingly, IAN did not only abolish the difference in lateral root density between Col-0 and *cyp79b2/b3* under salt stress, but additionally increased lateral root density for both genotypes in control conditions (p < 0.05) (Figure 6B). This increase in lateral root density in control conditions is caused by both a reduction in main root length and an increase in lateral root number (Supplemental figure 5). Because IAN is produced in emerging lateral roots during salt stress and can complement both lateral root phenotypes of the *cyp79b2/b3* mutant, we conclude that a lack of IAN could be responsible for the observed *cyp79b2/b3* root phenotypes.

**Figure 6:**
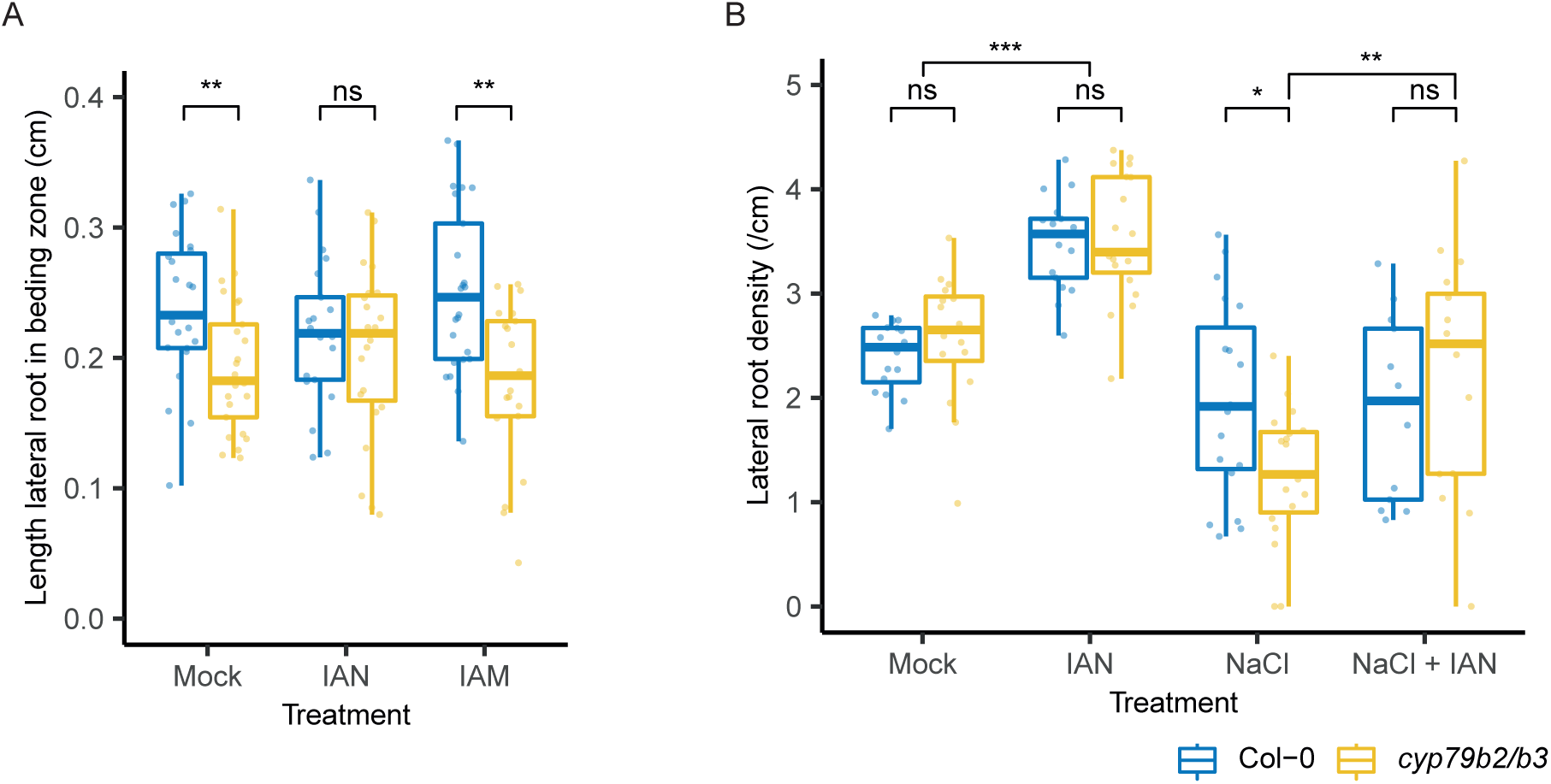
Exogeneous application of IAN complements the lateral root phenotypes of *cyp79b2/b3* mutants. A) Length of the lateral root in the bending zone of gravistimulated seedlings at 4 days after turning treated with mock, 5 µM IAN or 5 µM IAM. Statistical differences are indicated with brackets (two-way Anova, emmeans post-hoc test for genotypes per treatment, n > = 21). B) Lateral root density for 12 day-old seedlings, that were transferred to 0.5MS plates containing mock, 5 µM IAN, 125 mM NaCl or 5 µM IAN + 125 mM NaCl at 5 days after sowing. Brackets indicate statistical differences (pairwise Wilcoxon test, n > = 13), * p < 0.05, ** p < 0.01, *** p < 0.001.

## Discussion

We show here that *cyp79b2/b3* double KO mutant plants are not affected in their lateral root development or lateral root elongation after appearance but show a reduced lateral length just after emergence. Thereby, we identify CYP79B2/B3 as novel regulators of early lateral root elongation, which influences overall root architecture under salt stress. Despite the extensive knowledge about the processes that influence lateral root primordia initiation, development, outgrowth and emergence [reviewed in (Péret et al., 2009; Du and Scheres, 2018)], less is known about the formation of a new meristem in the developing lateral root and especially the activation of this meristem [reviewed in (Trinh et al., 2018)]. Marker gene expression patterns suggest that the formation of a new quiescent center in the lateral root primordia starts from stage 5 onwards (Goh et al., 2016), which coincides with the formation of an auxin maximum in the tip of the lateral root primordia (Benková et al., 2003). Besides having a quiescent center, an active meristem is constantly dividing and replenished [reviewed in (Trinh et al., 2018)]. A detailed report of cell divisions during lateral root formation showed that cell expansion is the main driver of lateral root emergence, while cell numbers only increase after emergence, proposing that the new lateral root meristem is only actively dividing after emergence (Malamy and Benfey, 1997). Furthermore, a recent report showed that mitotic quiescence of the quiescent center was only established after emergence, together with establishment of the cellular layout and unique gene expression profiles of the stem cell niche (Berthet et al., 2022). We found that the expression domain of *CYP79B3* shifts from a region underlying the lateral root primordia to the meristem of the lateral root after lateral root emergence. Interestingly, this shift does not coincide with the reported formation of a new auxin maximum or quiescent center identity at stage 5 (Benková et al., 2003; Goh et al., 2016), but only occurs after lateral root emergence. The role of CYP79B2 and CYP79B3 in lateral root growth most likely involves either maturation of the meristem after emergence or the promotion of lateral root growth, within the timeframe between lateral root emergence and macroscopic lateral root appearance.

Salt retards root development and reshapes root architecture (Julkowska et al., 2014). Depending on the exact conditions, different effects of salt on macroscopic lateral root density have been reported (Wang et al., 2009; Zolla et al., 2010; Julkowska et al., 2014), but clearly salt both reduces the initiation of lateral roots and the percentage of lateral roots that reach emergence (McLoughlin et al., 2012). Previously, *cyp79b2/b3* mutant roots were described to have both reduced lateral root density and lateral root length under salt stress but not control conditions (Julkowska et al., 2017). We now show that, after gravistimulation, the lateral root in the bending zone of *cyp79b2/b3* mutants is shorter compared to that of wild type plants in both salt and control conditions, while we also confirm the salt-dependence of the root architecture phenotype of 12-day old seedlings (Figure 6B). We cannot exclude that the early lateral root elongation phenotype in control conditions is caused by artificially forcing lateral root initiation through gravistimulation. However, another likely explanation for the root architecture phenotype only being apparent in salt stress is that the observed delay in lateral root appearance is relatively more important during the retarded root architecture development in salt stress. In this case, the higher rate of lateral root initiation (McLoughlin et al., 2012) and faster lateral root growth rate in control conditions could mask the small delay in lateral root appearance in *cyp79b2/b3* mutants. Furthermore, with respect to root architecture, CYP79B2/B3 were shown to influence lateral root density, but not consistently alter the underlying trait lateral root number during salt stress (Julkowska et al., 2017). The interaction between the delay in lateral root appearance and small temporal changes in main root growth could together result in the differential lateral root density phenotype of *cyp79b2/b3* in salt stress conditions. In conclusion, CYP79B2/B3 are important regulators of early lateral root elongation, which becomes apparent in macroscopic root architecture only under environmental stress conditions.

CYP79B2/B3 are responsible for converting tryptophane to iAOx at the start of the so-called iAOx biosynthesis pathway, which ultimately synthesizes various hormones and metabolites (Figure 1A). In the *cyp79b2/b3* mutant we found an upregulation of the genes encoding enzymes producing tryptophane and IG, which suggests the existence of a transcriptional feedback loop to upregulate the expression of upstream enzymes when there is a lack of iAOx or IGs. On the other hand, salt stress caused a downregulation of most IG producing enzymes. Furthermore, an upregulation of IAN producing enzymes and a salt induced, CYP79B2/B3-dependent IAN accumulation in emerging lateral roots was observed. Finally, exogenous application of IAN complemented the lateral root phenotypes of *cyp79b2/b3* for both root architecture and lateral root formation in a turning assay. Exogenous IAN application also increased lateral root density in Col-0 in control conditions. This phenotype can partly be explained by a reduced main root length in response to IAN (Supplemental figure 5), which is in accordance with previous reports (Lehmann et al., 2017). IAN also increases lateral root number in Col-0 in control conditions, but in the same conditions we do not see a further increase in lateral root length in response to a gravistimulus. This suggests that in control conditions, an excess amount of IAN influences lateral root initiation besides its effect on early elongation. Thus, our data suggests that IAN is one of the components in the iAOx pathway that alters root architecture under salt stress.

IAN can be converted to IAA and camalexin (Glawischnig et al., 2004; Nafisi et al., 2007; Böttcher et al., 2009; Sugawara et al., 2009). Recently, exogenous application of camalexin was shown to influence lateral root development, possibly influencing the transition from stage 2 to stage 3 (Serrano-Ron et al., 2021). However, in the *cyp79b2/b3* mutant we do not see an arrest in any of the stages during lateral root development. Moreover, we do not detect camalexin in our samples, making it less likely that the *cyp79b2/b3* phenotypes are caused by a decrease in camalexin. Auxin strongly steers lateral root development and growth [reviewed in (Du and Scheres, 2018)]. Our data show that salt decreases IAA levels in the bending zone of gravistimulated seedlings specifically in the early stages of lateral root development. This data is in line with previous studies that have shown that salt reduced auxin abundance and signaling in the primary root tip up to 1 day after treatment (Liu et al., 2015; Zhang et al., 2023). In contrast, salt has been shown to increase IAA levels in whole plants after 2 weeks of salt treatment (Cackett et al., 2022), showing how the effect of salt on auxin abundance could be dependent on the time of treatment or different between shoots and roots.

While salt increases IAN levels in a CYP79B2/B3-dependent manner and IAN can be converted to IAA, we did not find differences in IAA concentration in the *cyp79b2/b3* mutants compared to wild type in salt or control conditions. Interestingly, molecular docking simulations have shown that IAN can potentially bind the auxin receptor TRANSPORT INHIBITOR RESPONSE 1 (TIR1) (Vik et al., 2018). Furthermore, exogenous application of IAN increases the expression of *DR5::N7-VENUS,* a reporter of auxin responses, in the endodermis and the cortex of the main root (Katz et al., 2015). Thus, it possible that IAN influences auxin signaling without being converted to IAA by direct binding of IAN to the TIR1 receptor. On the other hand, we do not see an enrichment of auxin signaling genes within the DEGs of the *cyp79b2/b3* mutant compared to Col-0. Thus, even though we cannot exclude a local or transient change in IAA abundance, the observed changes in IAN do not coincide with changes in IAA, making it less likely that a reduced pool of IAA, converted from IAN, causes the delayed lateral root appearance in *cyp79b2/b3*.

In conclusion, we show that the iAOx pathway promotes early lateral root growth and that it contributes to regulation of lateral root density during salt stress. We provide evidence that supports a role for IAN as one of the metabolites explaining these phenotypes: salt increases the production of IAN and IAN complements root phenotypes of the *cyp79b2/b3* mutant. Further research could investigate its mode of action and potential cooperation with other metabolites in the iAOx pathway in stress physiology. Our work provides an important advance in understanding how the iAOx pathway contributes to root architecture plasticity under stress by influencing early lateral root elongation.

## Materials and methods

### Plant material and growth conditions

*Arabidopsis thaliana* wild-type Columbia-0 (Col-0) was used as a reference. The *cyp79b2-2/cyp79b3-2* knock-out mutant (Sugawara et al., 2009) was kindly provided by Hiroyushi Kasahara (RIKEN Center for Sustainable Resource Science, Yokohama, Japan). Plants were genotyped using the primers listed in supplemental table 1. Seeds were sterilized in 75 % EtOH and 25 % bleach for 5 min, washed twice in 70 % EtOH and once in 96 % EtOH, after which the seeds were left in the laminar flow to dry and were sown using 0.1 % agar on plates containing 0.5 x Murashige-Skoog (MS, Duchefa), 0.1 % M.E.S. Monohydrate (Duchefa) and 1 % Daishin agar (Duchefa) (pH 5.8, with KOH). Seeds were stratified at 4 °C for 2 days in the dark and grown at a 70° (root system architecture) or 90° (gravistimulation) angle in a growth chamber under long day conditions (16 h light: 8 h dark, 130 μmol/m^2^/s LED light) at 21 °C and 70% relative humidity. Seedlings were gravistimulated by turning 5 day-old seedlings 90° degrees.

### pCYP79B3::erCFP marker line construction

Gateway cloning was used to construct the *pCYP79B3::erCFP*. The 5.409 kb *CYP79B3* promotor was placed in front of erCFP (cyan fluorescent protein) and a NOPALINE SYNTHASE (NOS) terminator in the pGREENII-0125 vector. The primers used for amplifying the *CYP79B3* promotor are listed in supplemental table 2. The construct was transformed in Col-0 using Agrobacterium-mediated floral dip.

### Lateral root phenotyping of gravistimulated seedlings

Gravistimulated seedlings were transferred to plates containing 125 mM NaCl or control plates 3 h after turning. To determine time of appearance and lateral root elongation, seedlings were imaged using a timelapse imaging system with long day growth conditions (16 h light: 8 h dark, 130 μmol/m^2^/s LED light at 21 °C and 70% relative humidity). Plants were imaged every 20 min using an infrared camera. In the last image, the lateral root in the bending zone was traced using SmartRoot 2.0 (Lobet et al., 2011). The root length in previous images were corrected using an automated edge-detection script [as described in (Gigli-Bisceglia et al., 2022)]. The lateral root length was smoothed using a LOWESS smooth. A lateral root was categorized as appeared when the length > the average length of the first 60 hours + 0.01 cm. The microscopic length of emerged lateral roots was imaged using an EVOS FL Cell Imager. In this case, the length was determined from the tip of the emerged lateral roots to the stele of the primary roots, only roots that were positions perpendicular to the lens were included in the analysis. To determine the stage of lateral root primordia in the bending zone, seedlings were fixed [as described in (Voß et al., 2015)] at 1, 2, 3 and 4 days after turning and imaged using a Leica differential interference contrast optics microscope. The expression of *CYP79B3p::erCFP* lines was visualized in gravistimulated seedlings 2 and 3 days after turning, using a Leica TCS SP8 HyD confocal microscope, the CFP was excited with a 405 nm laser and measured from 463 nm to 549 nm.

### IAN complementation assays

For turning assays, five day-old seedlings were gravistimulated and immediately transferred to plates containing 5 µM IAN, 5 µM IAM or mock (0.005 % MeOH) treatment. Plates were scanned around 104 hours after turning and the length of the lateral root in the bending zone was measured. For root system architecture measurements, five or four day old seedlings were transferred to plates containing mock (0.005 % MeOH), 5 µM IAN, mock + 125 mM NaCl or 5 µM IAN + 125 mM NaCl. Roots were traced using SmartRoot 2.0 (Lobet et al., 2011).

### RNA isolation and sequencing

Seedlings were grown on plates covered with a mesh with 50 µm openings (Sefar BV). At 5 day-old, seedlings were gravistimulated and 3 h after turning transferred to plates containing 125 mM NaCl or control plates. A region of 3 mm around the bending zone was dissected, pooled for ∼40 seedlings per sample and frozen in liquid nitrogen. The samples were ground to powder using a paint shaker (∼10 Hz). From these samples, RNA was isolated using NZY Total RNA Isolation kit with on-column DNAse treatment (NZYtech). Quality control (Agilent Bioanalyzer 2100), RNA Poly-A enrichment library preparation and transcriptome sequencing were conducted by Novogene Co., LTD. Samples were sequenced on the NovaSeq 6000 platform in 150 bp paired-end mode.

### RNAseq analysis

Quality control, trimming and alignment of reads was performed using the nf-core pipeline [(Ewels et al., 2020), 10.5281/zenodo.1400710]. Based on the quality control and PCA plot (Supplemental figure 1A, B) sample 39 (Col-0, control, 2 days) was excluded from all analyses. Differential expression was determined using Wald tests for main and interaction effects in the DeSEQ2 package, the dataset was releveled on different timepoints or developmental stages. Analysis per timepoint and per developmental stage were loaded and performed separately. GO term analysis was performed using ClusterProfiler (Yu et al., 2012; Wu et al., 2021). Raw and processed data will be available upon publication at https://www.ebi.ac.uk/ with annotation number E-MTAB-13311.

### Hormone measurements

Hormone measurements were performed on similar plant material as used for RNA sequencing, grown as a separate replicate. Stable isotope-labeled internal standards (100 nM in 10% methanol) were added to ground samples (supplemental table 3, except for IAN). Extraction of hormones was performed as previously described (Floková et al., 2014), except a StrataX 30mg/3ml spe-column (Phenomenex) was used and solvents were removed using a speed vacuum system (ThermoSavant). Liquid chromatography-tandem mass spectroscopy was used for the detection and quantification of ABA, IAA, IAN and camalexin. Sample residues were dissolved in 100µL acetonitrile /water (20:80 v/v) and filtered using a 0.2 µm nylon centrifuge spin filter (BGB Analytik). A Waters XevoTQS mass spectrometer equipped with an electrospray ionization source coupled to an Acquity UPLC system (Waters) was used to quantify hormones by comparing retention times and mass transitions with standards as previously described (Schiessl et al., 2019; Gühl et al., 2021). Chromatographic separations were performed using acetonitrile/water (+ 0.1 % formic acid) on a Acquity UPLC BEH C18 column (2.1 mm x100 mm, 1.7µm, Waters) at 40 °C with a flowrate of 0.25 mL/min. The column was equilibrated for 30 minutes with the solvent (acetonitrile /water (20:80 v/v) + 0.1% formic acid). For analysis, 5 µL of sample was injected, followed by an elution program in which the acetonitrile fraction linearly increased from 20% (v/v) to 70% (v/v) in 17 minutes. Between samples, the acetonitrile fraction was increased to 100% and maintained there for one minute to wash the column. Before injecting the next sample, the acetonitrile fraction was set to 20 % in one minute and maintained at this concentration for one minute. A capillary voltage of 2.5 kV was used in combination with a source temperature of 150 °C and a desolvation temperature of 500 °C. Multiple reaction monitoring was used for quantification (Schiessl et al., 2019). The IntelliStart MS Console was used to optimize the MRM-settings for the different compounds (supplemental table 2). Peaks were analyzed using Targetlynx software and samples were normalized for the internal standard recovery (for IAA and ABA) and expressed relative to the sample fresh weight. A standard curve was used to convert peak area to pmol per mg fresh weight.

### Statistics

A Levene’s test and a Shapiro-Wilk test were used to test for equal variance and normal distribution. In case both tests were not significant, a two-way ANOVA was performed with an emmeans post-hoc test. Otherwise, a Kruskal Wallis test and a subsequent Wilcoxon test were performed. For timeseries data, a Benjamini-Hochberg adjustment was used to correct for multiple testing. For IAA, IAN and ABA measurements, the ARTool package (https://CRAN.R-project.org/package=ARTool) was used to perform a 3-way nonparametric factorial ANOVA (Wobbrock et al., 2011) with an ART-C (Elkin et al., 2021) post-hoc test. Finally, a chi-square test was used to compare the distribution of stages of lateral root primordia between Col-0 and *cyp79b2/b3* mutants. For all assays, seedlings that stopped growing after transfer, grew away from the plate, did not induce a lateral root in the bending zone or produced multiple lateral roots in the bending zone were excluded from the analysis.

## Acknowledgements

We thank Hiroyushi Kasahara (RIKEN Center for Sustainable Resource Science, Yokohama, Japan) for providing the *cyp79b2/b3* mutants. This work is supported by the European Research Council (ERC) under the EU Horizon 2020 Research and Innovation program (grant agreement 724321; Sense2SurviveSalt) to CT, the Netherlands Organization for Scientific Research (NWO) Veni (grant 863.15.010) to WK, the Netherlands Organization for Scientific Research (NWO) Vici (grant VI.C.192.033) to CT and the Netherlands Organization for Scientific Research (NWO) ALW Graduate Program (grant 831.15.004) to IK and CT.

## Author contributions

EvZ, IK, CT and CG planned and designed the research. EvZ, AJM, IK and KvdV carried out most experiments. TdZ, FV and WK optimized and carried out the hormone measurements. VW contributed the *pCYP79B3::erCFP* reporter line. EvZ and RH analyzed the data. EvZ, CT and CG wrote the manuscript. All authors read and provided feedback on the manuscript.

---

**Supplemental table 1:**
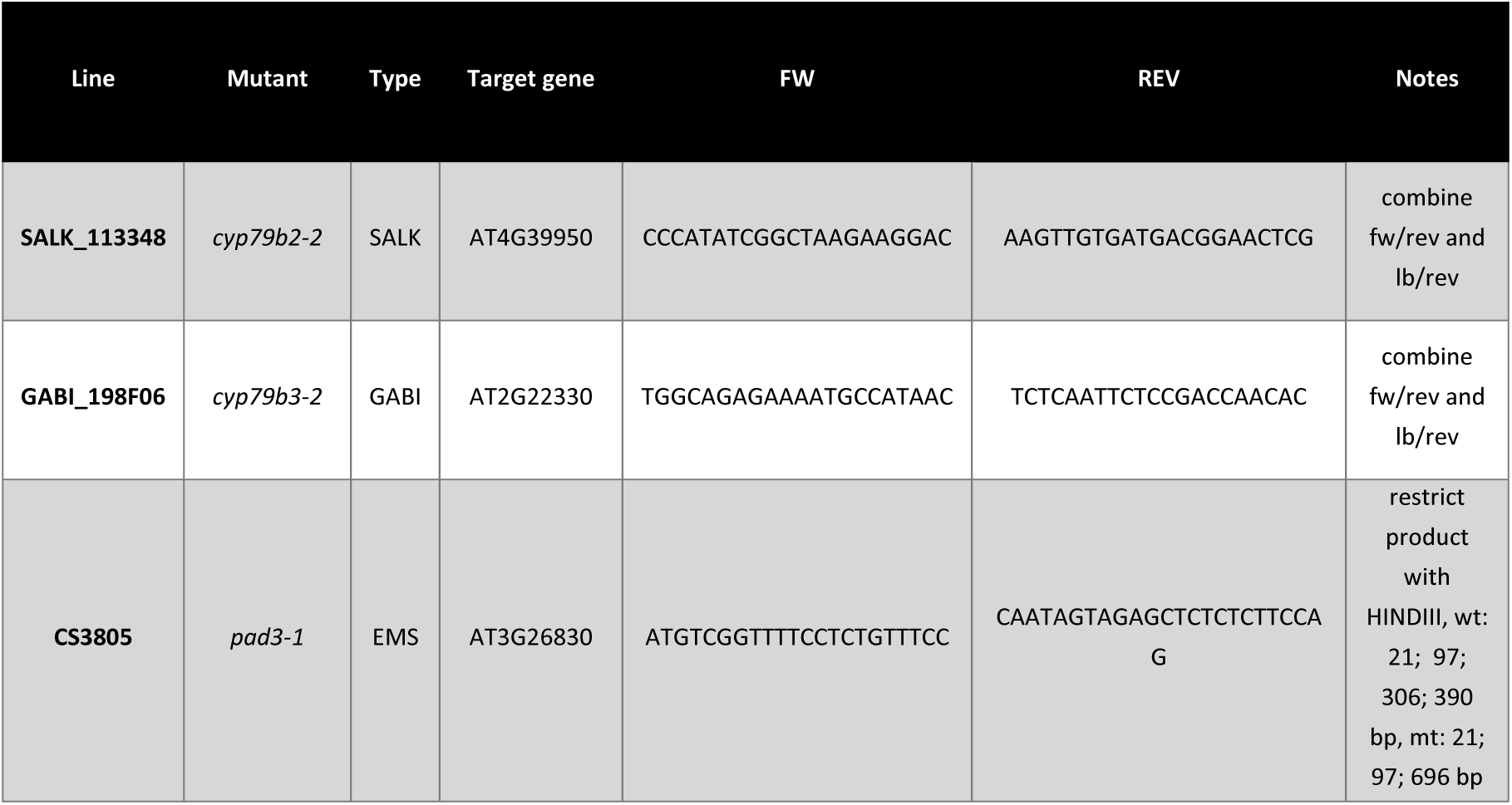
Genotyping primers used for validation mutants.

**Supplemental table 2:**
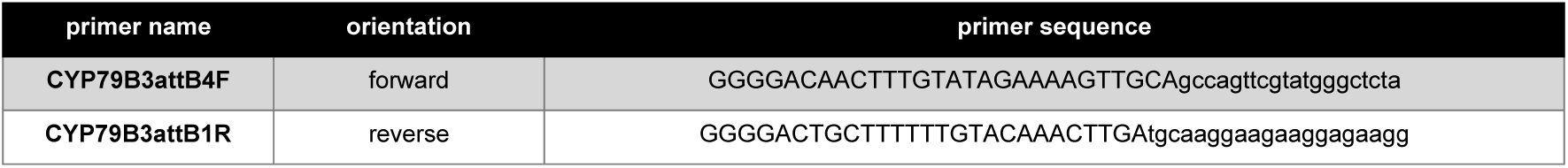
Primers used to amplify the CYP79B3 promotor.

**Supplemental table 3:** Multiple reaction monitoring (MRM) transitions for ABA, IAA, IAN and the used internal standards. *transition used for quantification.

| Compound | Retention time | Mass in negative ion mode | MRM transition | Cone V. | Coll. Energy |
| --- | --- | --- | --- | --- | --- |
| **ABA** | 4.15 | 263.25 | 153.15* | 30 | 10 |
|  |  |  | 219.15 | 30 | 15 |
| **[<sup>2</sup>H<sub>6</sub>]ABA** | 4.12 | 269.25 | 159.15* | 30 | 10 |
|  |  |  | 225.15 | 30 | 15 |
| **IAA** | 3.34 | 176.25 | 130.2* | 30 | 25 |
|  |  |  | 103.2 | 30 | 15 |
| **[<sup>13</sup>C<sub>6</sub>]IAA** | 3.32 | 182.1 | 136.2* | 30 | 25 |
|  |  |  | 109.2 | 30 | 15 |
| **IAN** | 5.44 | 157.2 | 130.05* | 30 | 30 |
|  |  | 157.2 | 117.0 | 30 | 20 |
|  |  | 157.2 | 90.0 | 30 | 10 |

**Supplemental figure 1:**
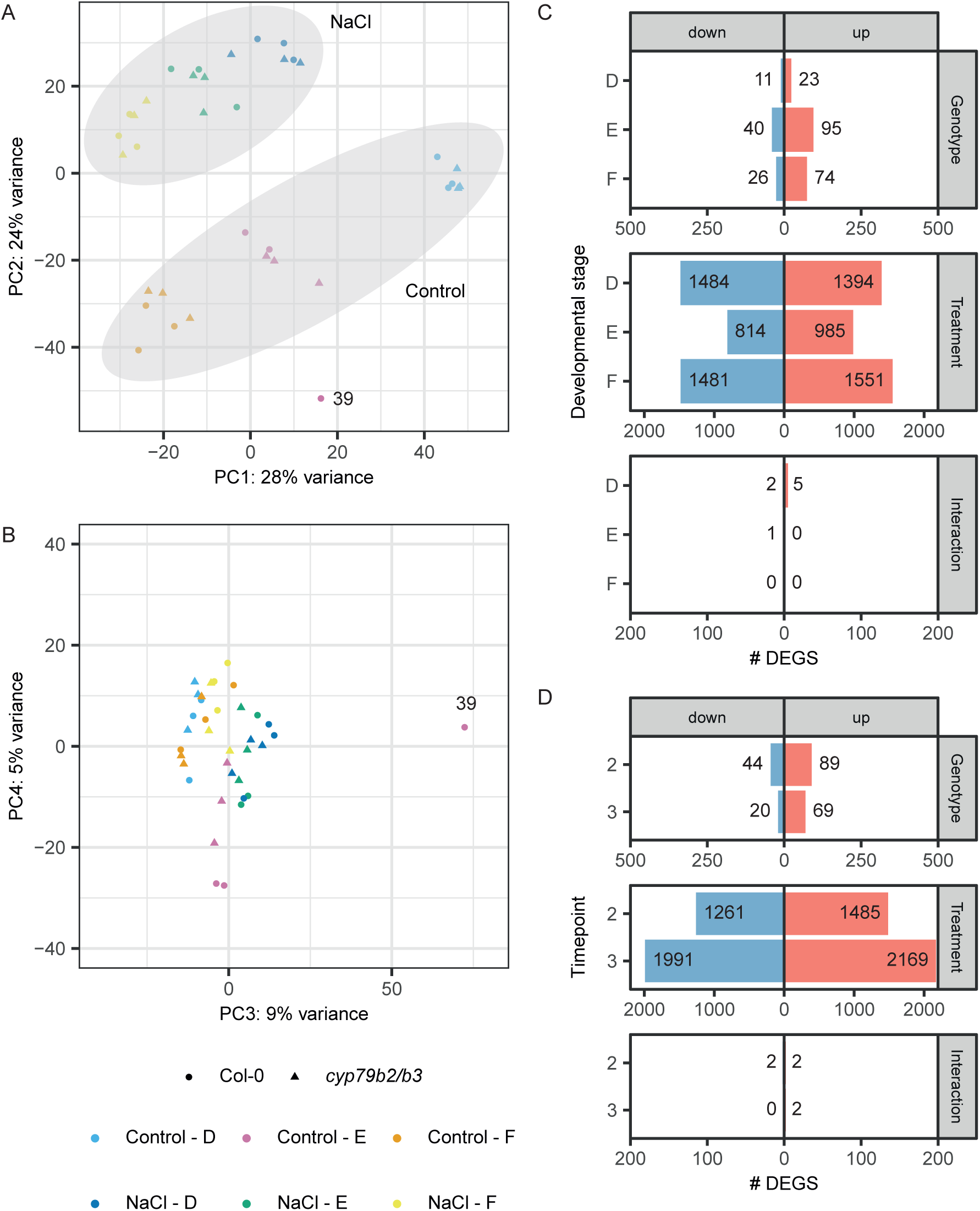
Number of DEGs and variation in RNAseq dataset. PCA of all samples in RNAseq dataset, showing A) PCA1 and PCA2 and B) PCA3 and PCA4. Sample 39 is marked (39) and identified as outlier. C-D) Number of DEGS (absolute LFC > 0.5, p < 0.05) for genotype effect (*cyp79b2/b3* vs Col-0), treatment effect (NaCl vs control) and interaction effect (effect of *cyp79b2/b3* mutation on the changes in expression during salt treatment). DEG analysis is performed C) per developmental stage and D) per timepoint after turning.

**Supplemental figure 2:**
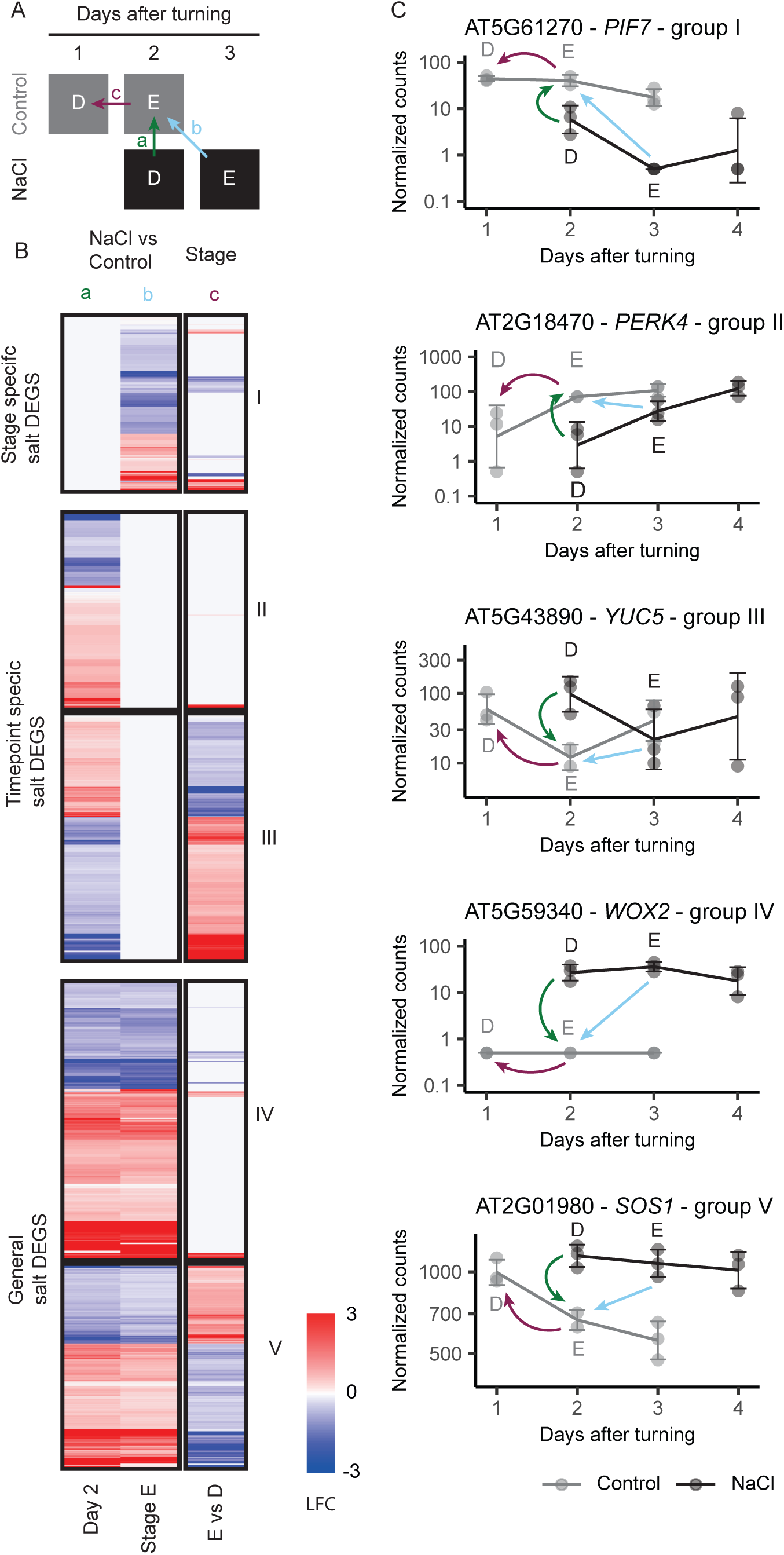
Differences between comparing gene expression per timepoint or per developmental stage at 2 days after turning. The bending zone of salt-treated gravistimulated seedlings was dissected and send for RNAseq. Salt-induced gene expressional during lateral root development and emergence is explored in Col-0. A) Scheme showing the between which samples the LFC was calculated to compare salt-induced gene expression at timepoint 2 (a) with salt-induced gene expression at stage E (b). The differential expression between stage E and D in control (c) was used to see how the developmental delay at timepoint 2 (stage E control, stage D salt) affected salt-induced gene expression at this timepoint (a). B) Heatmap showing LFC for effect a, b and c. Genes are placed in groups based on following criteria in 5 groups: stage specific salt DEGS (I, p < 0.05 for only b), timepoint specific DEGs (II & III, p < 0.05 for only a) and general salt DEGs (IV, V, p < 0.05 for a & b). Timepoint specific and general salt DEGs are subdivided in a group with (III & V) and without (II & IV) an opposite expression pattern for effect c (positive LFC for c effect and negative LFC for the a effect or the other way around). Only significant (p < 0.05) LFC are shown, non-significant LFC are in light grey. C) Normalized count for example genes in group I, III, IV and V in the bending zone of Col-0. Effect a, b and c are shown as arrows in the same colors as in A. Stage D and E are labelled. D = developing lateral roots, E = emerging lateral roots, F = fully emerged lateral roots.

**Supplemental figure 3:**
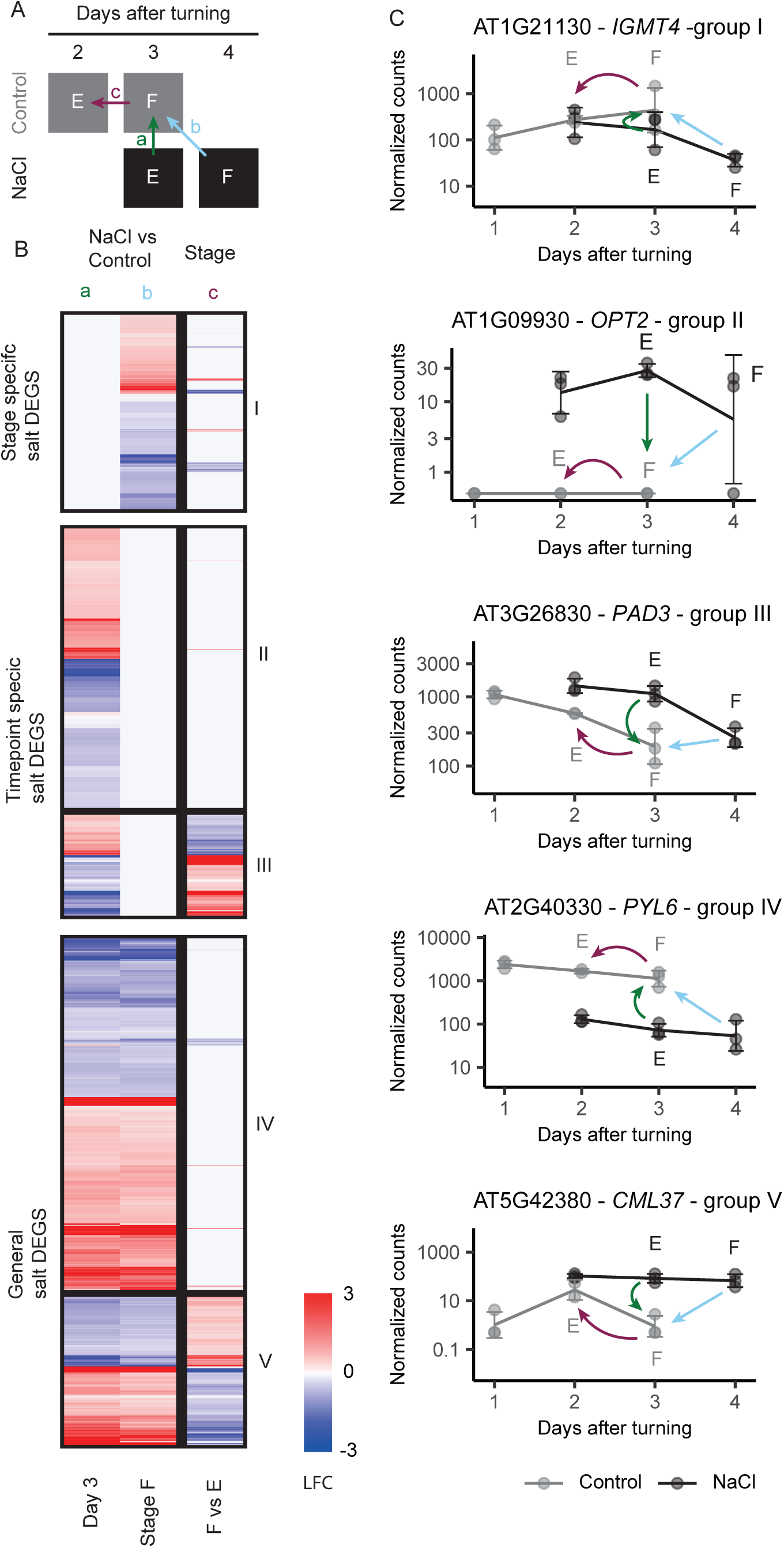
Differences between comparing gene expression per timepoint or per developmental stage at 3 days after turning. The bending zone of salt-treated gravistimulated seedlings was dissected and send for RNAseq. Salt-induced gene expressional during lateral root development and emergence is explored in Col-0. A) Scheme showing the between which samples LFC was calculated to compare salt-induced gene expression at timepoint 3 (a) with salt-induced gene expression at stage F (b). The differential expression between stage F and E in control (c) was used to see how the developmental delay at timepoint 3 (stage F control, stage E salt) affected salt-induced gene expression at this timepoint (a). B) Heatmap showing LFC for effect a, b and c. Genes are divided in 5 groups: stage specific salt DEGS (I, p < 0.05 for only b), timepoint specific DEGs (II & III, p < 0.05 for only a) and general salt DEGs (IV, V, p < 0.05 for a & b). Timepoint specific and general salt DEGs are subdivided in a group with (III & V) and without (II & IV) an opposite expression pattern for effect c (positive LFC for c effect and negative LFC for the a effect or the other way around). Only significant (p < 0.05) LFC are shown, non-significant LFC are in light grey. C) Normalized count for example genes in group I, III, IV and V in the bending zone of col-0. Effect a, b and c are shown as arrows in the same colors as in A. Stage D and E are labelled. D = developing lateral roots, E = emerging lateral roots, F = fully emerged lateral roots.

**Supplemental figure 4:**
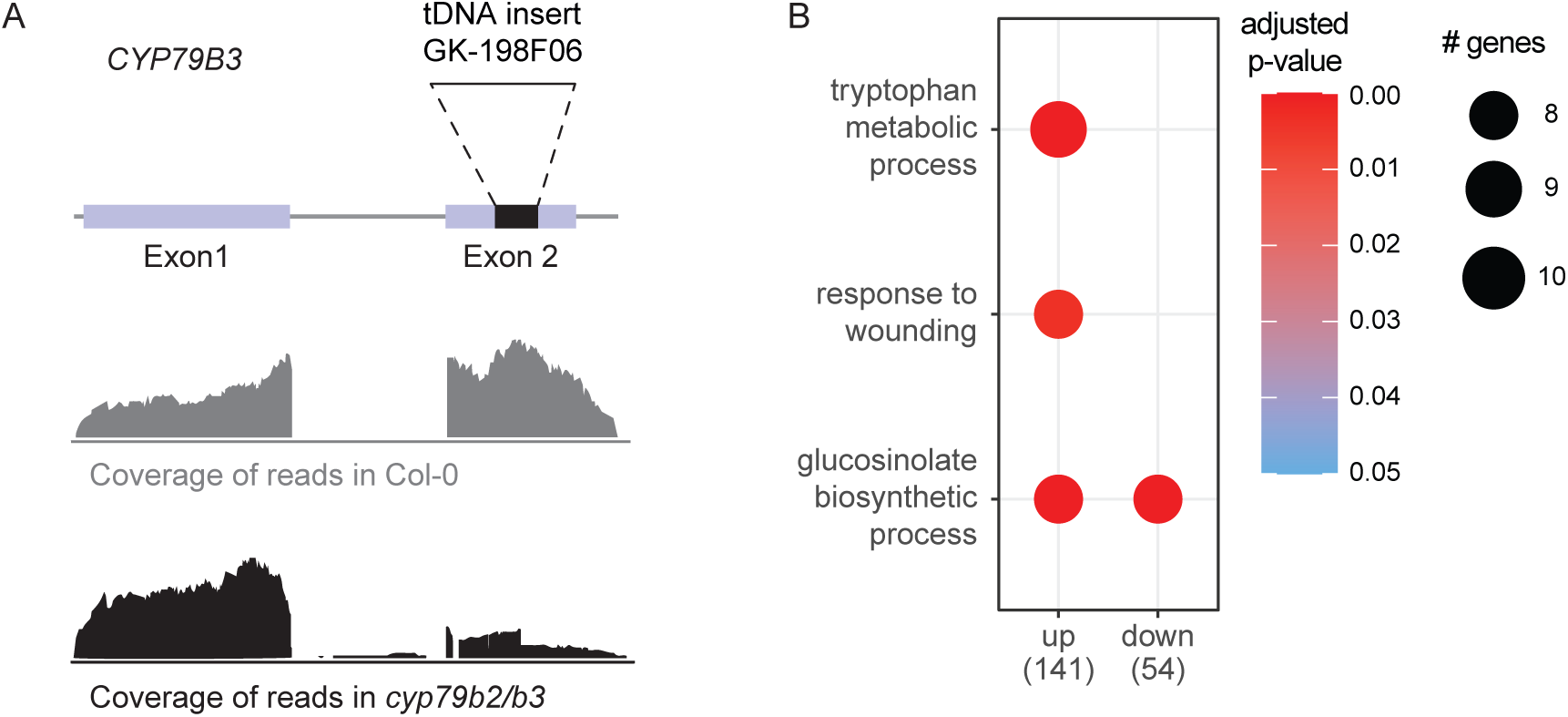
GO terms of genes with a differential expression in *cyp79b2/b3.* A) GO term enrichment of DEGS that were up or downregulated (p < 0.05 and absolute LFC > 0.5) in *cyp79b2/b3* compared to Col-0 at any developmental stage under control treatment. The significant enriched GO terms (p < 0.05) with more than 4 genes are shown. GO terms with more than 50% overlap were combined. Dot size represents the number of genes belonging to the term. Dot color represents the adjusted p-value. B) Read coverage on the *CYP79B3* gene in example samples of Col-0 and *cyp79b2/b3.* The location of the exons and t-DNA insertion are visualized.

**Supplemental figure 5:**
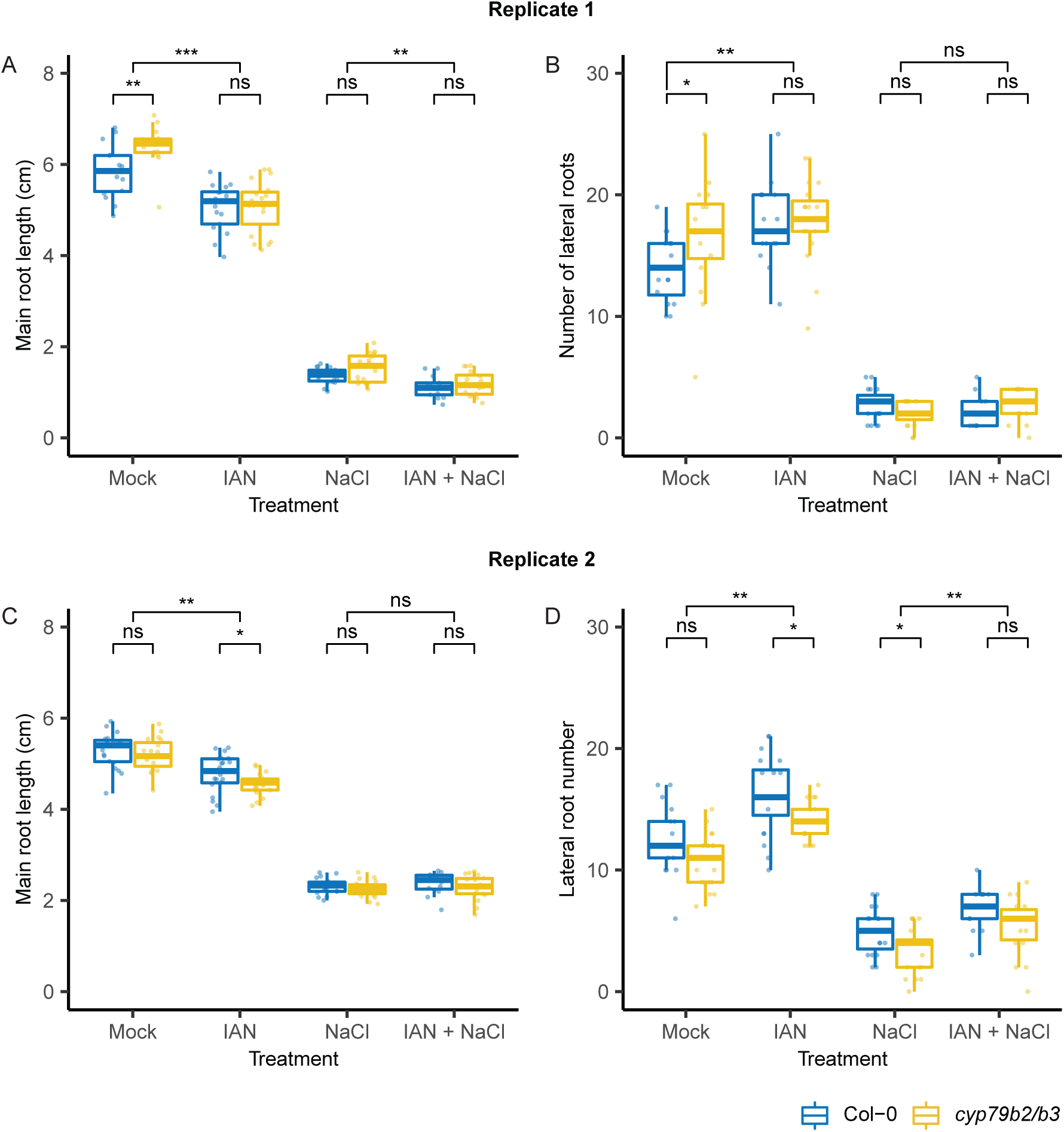
IAN reduces main root length and increases lateral root number in control conditions in Col-0. Seedlings were transferred to 0.5MS plates containing mock, 5 µM IAN, 125 mM NaCl or 5 µM IAN + 125 mM NaCl at 4 or 5 days after sowing. A, C) Main root length and B, D) lateral root number for 2 different replicates. Replicate 1 was traced at 12 day-old, replicate 2 was traced at 10 day-old to have similar sized root architectures. Brackets indicate statistical differences (pairwise Wilcoxon test, p < 0.05, n > = 13).

